# The role of *UBE3A* in the autism and epilepsy-related Dup15q syndrome using patient-derived, CRISPR-corrected neurons

**DOI:** 10.1101/2022.03.11.483963

**Authors:** Marwa Elamin, Aurelie Dumarchey, Christopher Stoddard, Tiwanna M. Robinson, Christopher Cowie, Dea Gorka, Stormy J. Chamberlain, Eric S. Levine

**Affiliations:** Neuroscience, University of Connecticut School of Medicine, Farmington, CT, USA; Genetics and Genome Sciences, University of Connecticut School of Medicine, Farmington, CT, USA

## Abstract

Chromosome 15q11-q13 duplication syndrome (Dup15q) is a neurodevelopmental disorder caused by maternal duplications of this region. Autism and epilepsy are key features of Dup15q, but affected individuals also exhibit intellectual disability and developmental delay. *UBE3A*, the gene encoding the ubiquitin protein ligase E3A, is likely a major driver of Dup15q because individuals with maternal, but not paternal 15q duplications have the disorder, and *UBE3A* is the only imprinted gene expressed solely from the maternal allele in mature neurons. Nevertheless, the exact role of *UBE3A* is yet to be determined. To establish whether *UBE3A* overexpression is required for Dup15q neuronal deficits, *UBE3A* expression was manipulated in patient-derived induced pluripotent stem cell (iPSC) lines using antisense oligonucleotides. Dup15q neurons exhibited hyperexcitability features compared to genome-edited isogenic control neurons, and this phenotype was generally prevented by normalizing *UBE3A* levels throughout *in vitro* development. Overexpression of *UBE3A* in an iPSC line with a paternal duplication resulted in a profile similar to that of Dup15q neurons except for synaptic phenotypes. These results indicate that *UBE3A* overexpression is necessary for most Dup15q cellular phenotypes. However, the inability of *UBE3A* overexpression to recapitulate synaptic phenotypes suggests an important role for non-imprinted genes in the disorder.

## Introduction

Chromosome 15q11-q13 duplication syndrome (Dup15q) is a neurodevelopmental disorder caused by duplications of the 11.2-13.1 region within the long arm of chromosome 15. There are two major genetic subtypes of Dup15q: idic(15), which is caused by an isodicentric supernumerary chromosome that carries two extra copies of the 15q11.2-q13.1 region, and int(15), which arises from a maternal interstitial duplication of the same region. In addition to these two major subtypes, some individuals have a maternal interstitial triplication of 15q11.2-13.1. Autism and seizures are two of the most common behavioral phenotypes of the syndrome (1). Individuals with Dup15q also present with intellectual disability and substantial fine and gross motor deficits. Clinical studies report that 77 - 100% of Dup15q patients are affected by autism (2, 3), and 63% of individuals with idic(15) experience seizures, which often present with multiple types and require aggressive treatment with broad-spectrum antiepileptic drugs (4). Moreover, epileptic children with Dup15q have an increased risk of Sudden Unexpected Death in Epilepsy (SUDEP) (5), therefore, understanding the cellular and molecular mechanisms that lead to seizure generation is of great importance.

In general, two extra maternal copies of 15q11-q13, carried either on an isodicentric extra chromosome or as part of an interstitial triplication, are associated with more severe behavioral, cognitive, and seizures phenotypes compared to maternal interstitial duplications, which have one extra copy of the genetic region (6). The extra genetic material in Dup15q includes approximately 20 genes (7), and active research is devoted toward understanding and identifying the genes that contribute to phenotypes associated with the disorder. Candidate genes include *UBE3A*, which encodes an ubiquitin ligase, *GABRA5, GABRB3*, and *GABRG3*, which encode subunits of the GABA_A_ receptor, *ATP10*, which encodes a mitochondrial inner membrane protein, *HERC2*, a ubiquitin ligase known to interact with and potentiate UBE3A activity, and others (7).

Overexpression of *UBE3A* is likely a major driver of Dup15q phenotypes. *UBE3A* encodes an E3 ubiquitin ligase that targets proteins for degradation by the proteasome (8). Moreover, it has been reported to act as a transcriptional coactivator for steroid receptors (9), and it has been implicated in the regulation of multiple genes associated with nervous system development (10). *UBE3A* is the only gene in the region expressed solely from the maternal allele, as *UBE3A* is paternally imprinted and silenced in mature neurons; hence, a maternal duplication would increase its gene dosage while a paternal duplication would not (11). Dup15q is fully penetrant in individuals with maternal duplications, while individuals with paternal duplications are usually unaffected or mildly affected (3, 12), supporting a key role for *UBE3A*. Nevertheless, the contribution of other duplicated genes in the region to Dup15q phenotypes is unknown.

The generation of animal models for Dup15q has been difficult. Creation of an interstitial duplication of the syntenic region in mice resulted in a mouse model with excellent construct validity (13). However, this model has poor face validity as there were no behavioral or physiological phenotypes with maternal duplication, whereas paternal duplication resulted in subtle autism-like phenotypes (13). Overexpression of *UBE3A* alone resulted in mice with either mild autistic-like behaviors (14, 15) or minor changes in seizure susceptibility (16). However, these models are not ideal for recapitulating *UBE3A* genetic overdose because the transgenes encoded only one of three potential protein isoforms and were epitope-tagged, potentially disrupting the ubiquitin ligase function of UBE3A. The failure to encompass the full range of Dup15q phenotypes in animal models and replicate the complex genetics of the disease makes the use of induced pluripotent stem cell (iPSC)-derived neurons from Dup15q patients an excellent choice to study the syndrome.

The goal of this study was to determine the role of *UBE3A* overexpression in Dup15q neuronal phenotypes. We used an innovative approach to generate an isogenic control line for an idic(15) patient-derived iPSC line. Using these lines, we normalized *UBE3A* levels with ASOs in Dup15q neurons to determine whether *UBE3A* overexpression is necessary for establishing cellular phenotypes. To explore whether *UBE3A* overexpression was sufficient to cause Dup15q cellular phenotypes, we used ASOs to increase *UBE3A* expression in neurons derived from an individual with a paternal interstitial duplication.

## Results

### Elimination of the isodicentric chromosome 15

We previously showed that patient-derived iPSCs carry the exact genetic makeup of individuals affected by Dup15q and maintain their methylation imprint following the reprogramming process (7). In a subsequent study (comparing multiple patient-derived Dup15q and control iPSC lines), we uncovered several hyperexcitability phenotypes in Dup15q neurons, including increased excitatory synaptic event frequency and amplitude and increased action potential (AP) firing (17). Although comparisons that involve unrelated control and disease lines provide important and relevant information, inherent functional differences between cell lines (which are likely driven by different genetic backgrounds), can obscure important cellular phenotypes. The use of isogenic cell lines provides a better approach to investigate cellular deficits in disease models, and the development of the CRISPR/Cas9 gene-editing technology has greatly enabled this process, especially for monogenic diseases (18-20).

We sought to generate an isogenic control for a Dup15q idic(15) patient-derived iPSC line to more confidently ascribe Dup15q cellular phenotypes to the extra genetic material on the idic(15) chromosome. We employed a strategy previously used to eliminate chromosome 21 in a trisomy 21 cellular model and the Y chromosome in typical male iPSCs (21, 22). This strategy entailed the use of CRISPRs to cut multiple times within chromosome 15q-specific repeats. We nucleofected Dup15q iPSCs with a cocktail of three different CRISPRs targeting *GOLGA8, SNORD116*, and *SNORD115* (Supplemental Table 1). The guide RNAs designed to target these loci are each predicted to cut multiple times within chromosome 15q11.2-q13.3 region but are not found outside of the chromosome. Our rationale was that these CRISPRs would make multiple double-stranded breaks on the two chromosomes 15 as well as the supernumerary idic(15) in the iPSC line. Since the idic(15) has two copies of 15q11.2-q13.3 and very little additional sequence, we reasoned that the entire chromosome might be lost upon becoming “shredded” by the CRISPR cocktail. Following transient selection for the CRISPRs, clones were screened using quantitative copy number assays for *UBE3A* and *PML*. Idic(15) iPSCs have four copies of *UBE3A*, but only two copies of PML, which is located near the distal end of chromosome 15. Four out of 58 clones showed a copy number decrease in *UBE3A* with no decrease in PML copy number.

The four clones with reduced *UBE3A* dosage were expanded and subjected to quantitative DNA methylation analysis at the *SNRPN* locus. Clones that had lost the idic(15) chromosome would have 50% methylation at *SNRPN* due to the presence of one maternal and one paternal chromosome 15. Deletions that included the imprinted domain of maternal chromosome 15 or paternal chromosome 15 would have 66% and 100% methylation at *SNRPN*, respectively. Following this screening paradigm, one single clone from a starting pool of 58 was identified with the appropriate copy number and DNA methylation parameters (Fig. 1A). CytoSNP and karyotype analysis confirmed the loss of the idic(15) chromosome without any other detectable copy number changes or structural rearrangements (Fig. 1B, Supplemental Table 2). The presence of non-chromosome 15 copy number changes in the edited iPSCs that were also present in the idic-1 iPSC line confirmed that the cytoSNP profiles and karyotypes of these two cell lines were identical to each other, except for the 15q copy number changes and supernumerary chromosome (Supplemental Table 2). Finally, RT-qPCR in normal, idic-1, and isogenic corrected control iPSCs demonstrated that gene expression of both imprinted and non-imprinted genes in the 15q region was reduced in the corrected control line compared to idic-1 and very similar to expression levels in unaffected control iPSCs (Fig. 1C).

**Figure 1.**
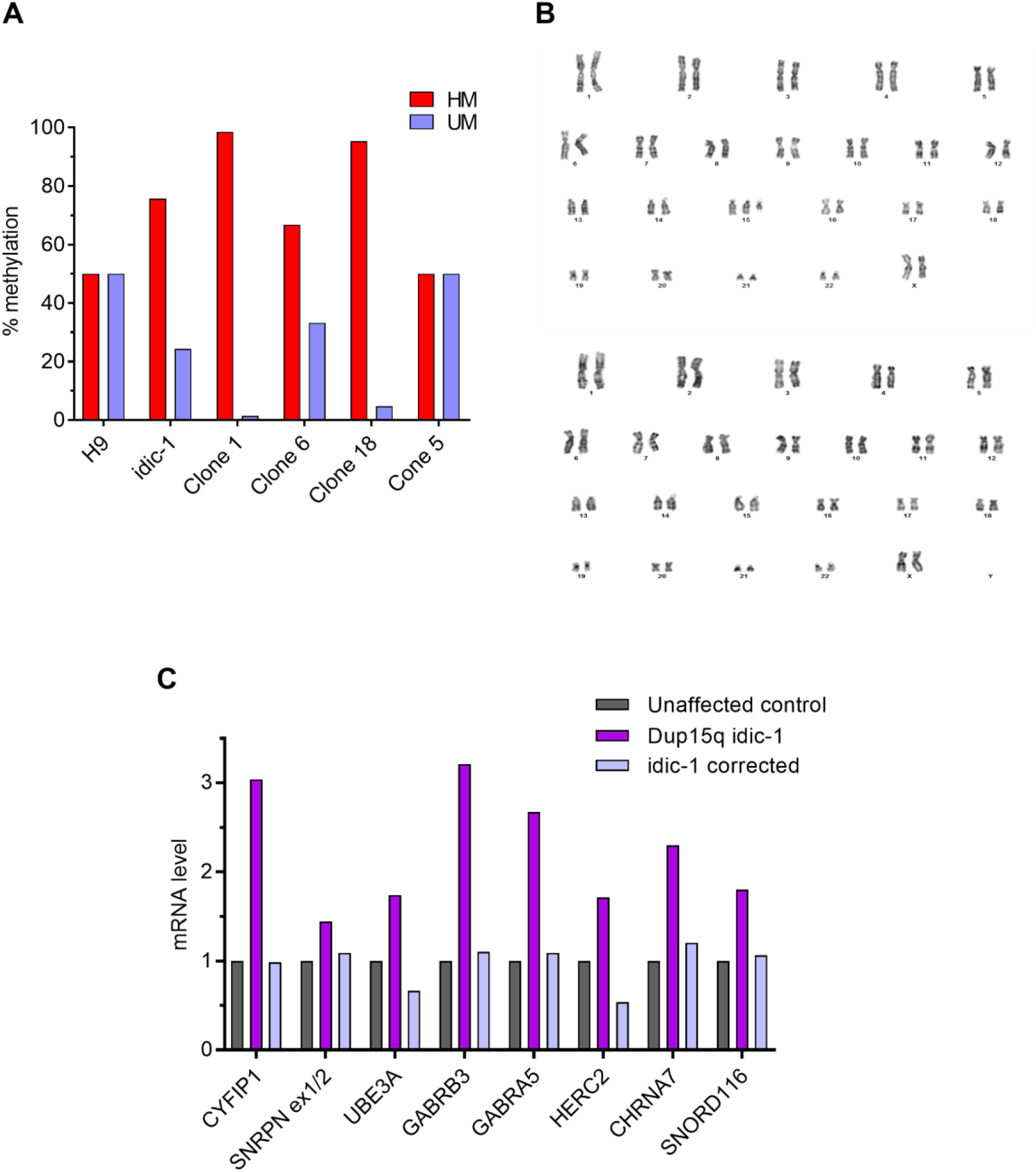
Generation of Dup15q isogenic corrected iPSC line. **(A)** Methylation status of clones compared to idic-1 Dup15q parent line and H9 control line. HM: hypermethylated, UM: unmethylated. **(B)** Karyotyping of idic-1 iPSC line (top) and corrected clone 5 of idic-1 (bottom). Note the loss of the idic(15) chromosome. **(C)** qRT-PCR analysis of imprinted and non-imprinted gene expression in the 15q locus in idic-1 iPSCs and the corrected idic-1 line. RNA levels are presented relative to an unaffected control iPSC line.

### Hyperexcitability of Dup15q neurons during *in vitro* development

To characterize the electrophysiological properties of Dup15q neurons, the idic-1 cell line and its CRISPR-corrected isogenic counterpart were differentiated into neurons using a modified dual-SMAD inhibition protocol (Supplemental Fig. 1A). Differentiation via this protocol yielded neuronal cultures that consisted of MAP2-positive neurons expressing the glutamatergic marker TBR1 (70 - 80%) and GABAergic neurons expressing GAD65, as well as astrocytes expressing S100β (7, 17).

Whole-cell patch clamp recordings were conducted at 8 weeks and 19 weeks of *in vitro* development. Dup15q neurons showed a significantly greater inward sodium current density compared to the corrected line at both time points, whereas the transient outward potassium current density was significantly increased at 19 weeks (Fig. 2A). Consistent with the increased inward and outward currents, Dup15q neurons had an increased AP amplitude and decreased AP width (Fig. 2B). Increased frequency of induced AP firing and a hyperpolarized AP threshold were also observed in Dup15q neurons at both time points (Fig. 2C). Dup15q neurons showed normal maturation of the resting membrane potential (RMP) compared to their corrected counterparts and no difference in cell capacitance (Supplemental Fig. 1B, C). Input resistance was decreased in Dup15q neurons at both time points (Supplemental Fig. 1D).

**Figure 2.**
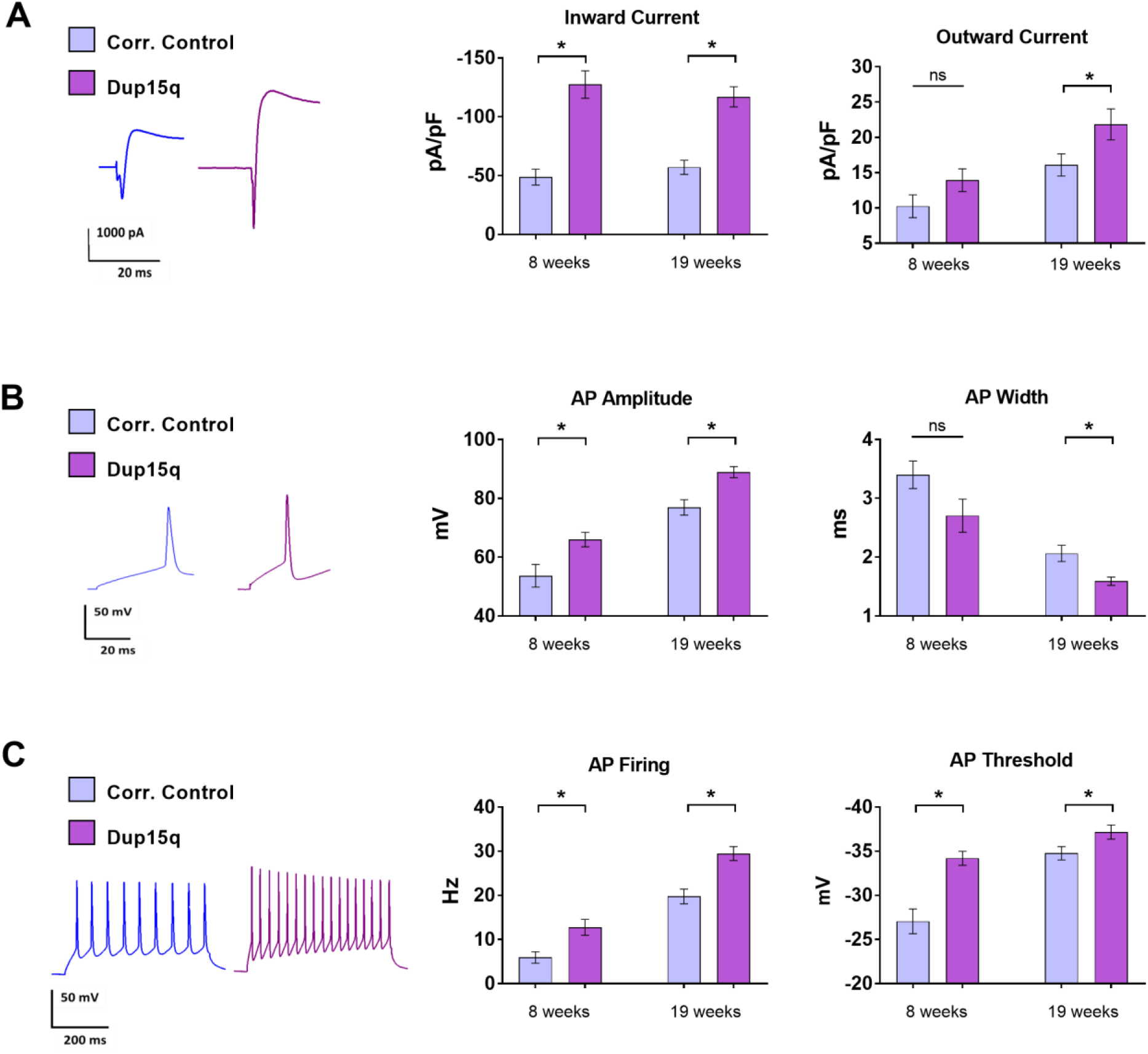
Cellular phenotypes of idic(15) Dup15q neurons compared to isogenic corrected control. **(A) Left**: Example traces of inward and outward currents elicited with a voltage step from -70 mV to +40 mV at 19 weeks of *in vitro* development. **Middle:** Maximum inward current density. **Right:** Maximum outward current density. **(B) Left:** Example action potential (AP) traces elicited by +80 pA current step. **Middle:** Peak AP amplitude. **Right:** AP width at half-maximal amplitude **(C) Left:** AP traces during a 500 ms current step elicited by +80 pA. **Middle**: Maximum AP firing rate. **Right:** AP threshold. n = 22 - 25 cells per group. Student’s t-test; *, p< 0.05.

### Altered synaptic transmission and spontaneous firing in Dup15q neurons

Voltage-clamp recordings of Dup15q neurons and corrected isogenic controls revealed a significant increase in the frequency and amplitude of spontaneous excitatory postsynaptic currents at 19 weeks (sEPSCs; Fig. 3A), similar to our previous study comparing Dup15q lines to unaffected controls (17). There was also a decrease in the frequency, but not amplitude, of spontaneous inhibitory postsynaptic currents (sIPSCs; Fig. 3B). Spontaneous postsynaptic currents consisted of both miniature postsynaptic currents (mPSCs) and AP-dependent currents. To isolate AP-independent miniature synaptic events, a separate series of recordings were carried out in the presence of 1 μM tetrodotoxin (TTX). Dup15q neurons showed an increase in mEPSC frequency and amplitude (Fig. 3C). There were no significant differences in mIPSCs (Fig. 3D). The increase in mEPSC frequency may reflect increased synapse number and/or increased presynaptic release probability. Immunostaining for the postsynaptic marker PSD95, however, failed to detect a statistically significant increase in PSD95 puncta in Dup15q neurons (Fig. 3E). The increased sEPSC amplitude may also reflect an increase in spontaneous neuronal firing since AP-dependent events are associated with higher amplitudes compared to miniature synaptic events. Supporting this, there was a significant increase in the number of AP-dependent calcium transients in Dup15q neurons (Fig. 3F)

**Figure 3.**
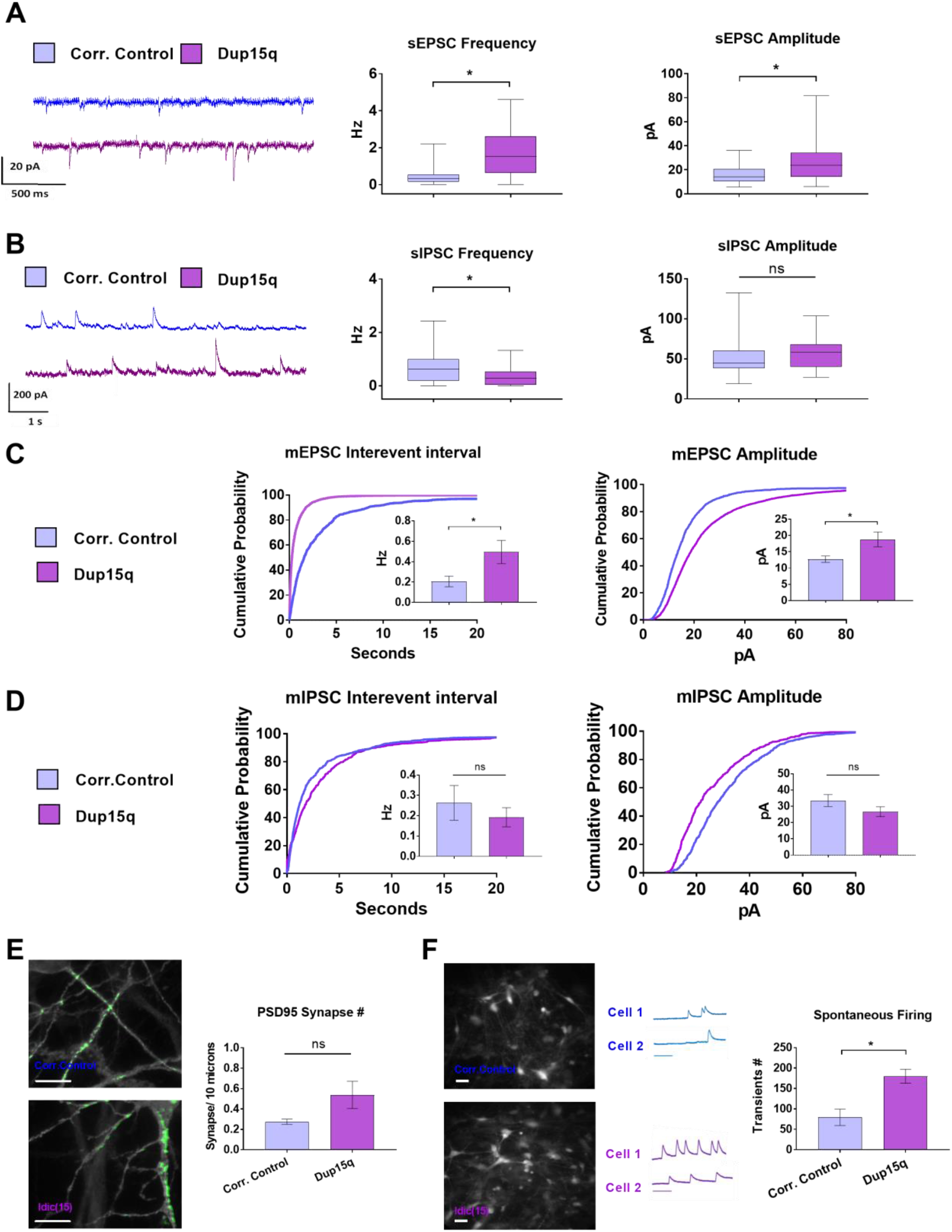
Synaptic transmission and spontaneous firing in Dup15q neurons. **(A) Left:** Example sweeps of spontaneous excitatory postsynaptic currents (sEPSCs) at 19 weeks. Cells were voltage-clamped at -70 mV. **Middle:** sEPSC frequency at 19 weeks. **Right:** sEPSC amplitude at 19 weeks. Box and whisker blots represent the first quartile, median, and third quartile; bars represent 2.5-97.5 percentile outliers (n = 22 -25 cells per group). Mann Whitney U test; *, p< 0.05. **(B) Left:** Example sweeps of spontaneous inhibitory postsynaptic currents (sIPSCs) at 19 weeks. Cells were held at 0 mV. **Middle:** sIPSC frequency at 19 weeks. **Right:** sIPSC amplitude at 19 weeks. Box and whisker blots as above (n = 22 -25 cells per group). **(C)** Miniature excitatory postsynaptic currents (mEPSCs) recorded at -70 mV in the presence of 1 μM TTX. **Left:** Cumulative probability histogram of the interevent interval of all mEPSC events. **Inset**: Frequency of mEPSC per cell, represented as mean +/- SEM. **Right:** Cumulative probability histogram of the amplitude of all mEPSC events. **Inset**: Amplitude of mEPSC per cell, represented as mean +/- SEM (n>20 cells per group). Unpaired t-tests; *, p< 0.05. **(D)** Miniature inhibitory postsynaptic currents (mIPSCs) recorded at -70 mV in the presence of 1 μM TTX. **Left:** Interevent interval of all mEPSC events recorded across all cells. **Inset**: Frequency of mIPSCs per cell. **Right:** Cumulative probability histogram of the amplitude of all mIPSC events recorded across all cells. **Inset**: Amplitude of mIPSC per cell (n= 15 cells in each group). Student’s t-test; *, p< 0.05. **(E)** Representative images of PSD95 immunostaining and density of PSD95 puncta in 19-week old neurons (n=4-5 coverslips per group, unpaired t-test). Scale bar: 10 μm. **(F)** Calcium imaging. **Left:** Representative images of Dup15q and corrected neurons after incubation with X-Rhod-1 fluorescent dye. Scale bar: 50 μm; **Middle**: Example traces of spontaneous calcium transients. Scale bar: 100 sec. **Right:** Number of spontaneous calcium transients per coverslip over 15 min. (n=5 coverslips per group). Unpaired t-test; *, p<0.05.

### The contribution of *UBE3A* overexpression to Dup15q cellular phenotypes

*UBE3A* is exclusively expressed from the maternal allele in mature neurons, thus overexpression of this gene is thought to be a critical factor in the development of the maternally-inherited syndrome. To examine the contribution of *UBE3A* to Dup15q cellular phenotypes, we normalized expression in idic(15) Dup15q neurons using ASOs targeting *UBE3A*. ASOs bind to RNA in a sequence-specific manner and recruit RNAse H to cleave the RNA bound to the DNA-like core of the ASO (23, 24). This, in turn, recruits exonucleases leading to the knockdown of the target RNA. Intriguingly, ASOs are freely taken up by neurons without the use of transfection reagents, allowing for the knockdown of the intended genes in every neuron within the culture (24). Furthermore, ASO effects are concentration-dependent and highly stable, with a single treatment leading to enduring knockdown in post-mitotic neurons (24). We have previously designed and characterized ASOs that selectively reduce the expression of *UBE3A* with high efficiency, resulting in immediate, and long-lasting, knockdown of both mRNA and protein levels (25). To normalize *UBE3A* levels in Dup15q neurons, ASO treatment (10 µM) was initiated at two time points: at 6 weeks *in vitro*, to identify a role of *UBE3A* in the establishment of cellular phenotypes, and at 16 weeks, to determine if established phenotypes can be reversed. Patch clamp recordings were performed at week 19 (Fig. 4A). Neurons were collected 2-3 weeks post-treatment and the level of *UBE3A* mRNA was confirmed via qPCR (Fig. 4B). A scrambled sequence ASO was used to control for non-specific effects.

**Figure 4.**
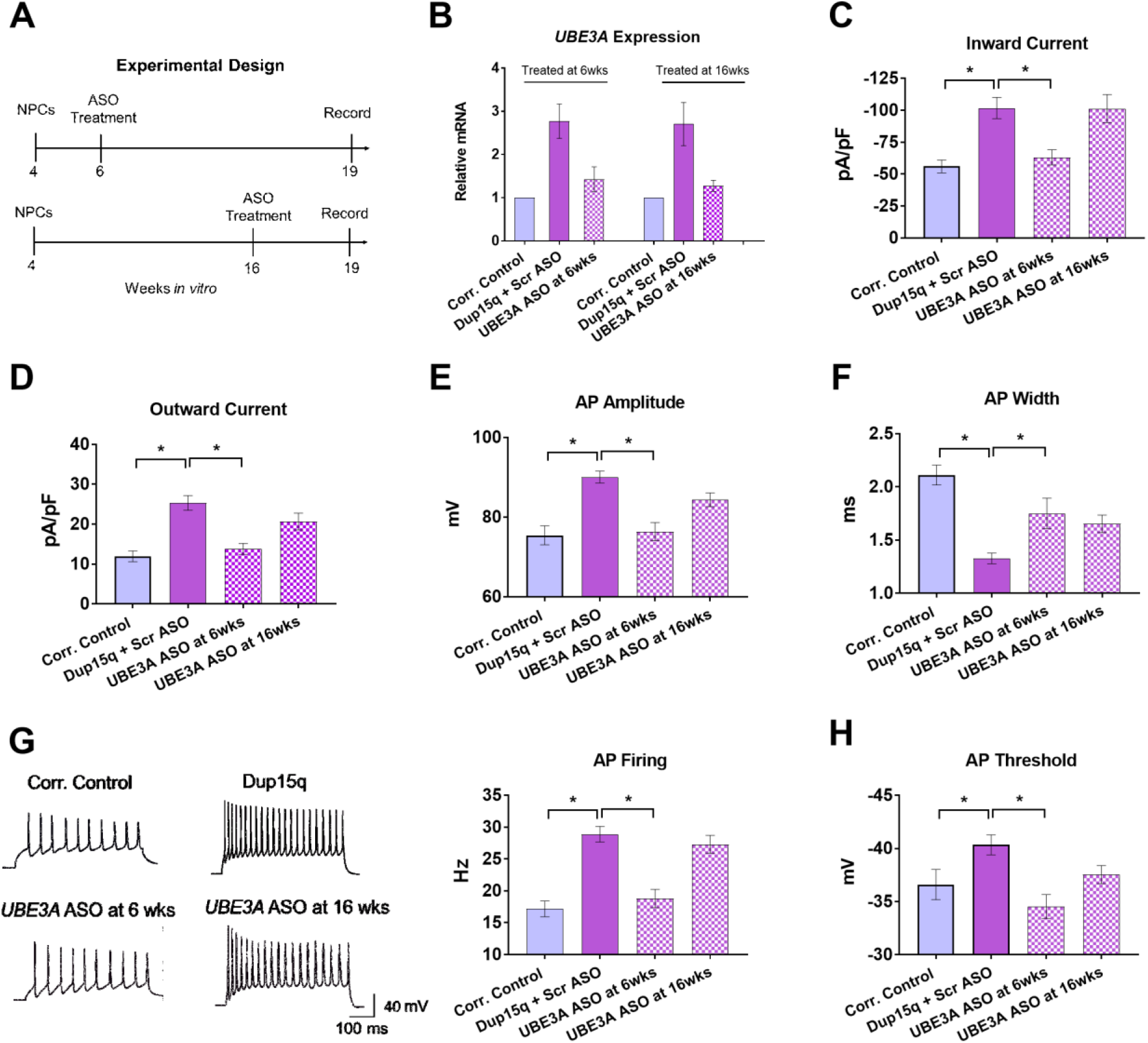
Normalization of *UBE3A* expression prevents intrinsic phenotypes in Dup15q neurons. **(A)** Experimental design of *UBE3A* ASO treatments and recording times. **(B)** Relative expression of *UBE3A* mRNA in corrected control neurons, Dup15q neurons + scramble (Scr) ASO, and Dup15q neurons + *UBE3A* ASO at 6 weeks and 16 weeks. PCR performed 2-3 weeks post-treatment **(C)** Maximum inward current density. **(D)** Maximum outward current density. (E) Action potential (AP) amplitude. **(F)** AP width. **(G)** Left: Representative traces of AP firing at 19 weeks during a 500 ms current step from -10 pA to +80 pA. Right: AP firing rate. **(H)** AP threshold. (n = 22-30 cells/group). One-way ANOVA and Dunnett’s multiple comparisons tests. *, p< 0.05.

Normalizing *UBE3A* expression at 6 weeks prevented the increased inward and outward currents (Fig. 4C, D), the increased AP amplitude (Fig. 4E), the decreased AP width (Fig. 4F), the increased AP firing rate (Fig. 4G), and the hyperpolarized AP threshold (Fig. 4H) when recordings were carried out at 19 weeks. Interestingly, *UBE3A* normalization at 16 weeks failed to reverse these phenotypes at 19 weeks (Fig. 4C - H). To determine if normalizing *UBE3A* at 6 weeks prevented the expression of these phenotypes within 2 weeks of treatment, we performed patch clamp recordings in 8 week-old neurons after treatment with *UBE3A* ASO at 6 weeks. The increase in inward current was completely normalized at 8 weeks (Supplemental Fig. 2A). A similar trend was seen in outward current, AP firing rate, amplitude, and threshold, although not statistically significant (Supplemental Fig. 2B - D). To determine if ASO treatment had non-specific effects on electrophysiological parameters, we compared scramble ASO-treated cells to untreated cells and found no differences between the two groups (Supplemental Fig. 3). To confirm the role of *UBE3A* in the aforementioned phenotypes, we normalized *UBE3A* levels in a different Dup15q line with the same genetic aneuploidy (idic-2). Normalization of *UBE3A* levels at 6 weeks decreased inward current, outward current, AP amplitude, and AP firing rate (Supplemental Fig. 4). In summary, *UBE3A* normalization at 6 weeks prevented the development of neuronal hyperexcitability, indicating that *UBE3A* overexpression is necessary for the development of these cellular phenotypes.

To investigate the contribution of *UBE3A* to the synaptic phenotypes in Dup15q neurons, we used ASOs to normalize *UBE3A* levels starting at either 6 or 16 weeks of *in vitro* development and recorded synaptic activity at 19 weeks. *UBE3A* normalization at 6 weeks prevented the increased sEPSC frequency and amplitude in Dup15q neurons, while treatment at 16 weeks failed to do so (Fig. 5A). Interestingly, normalization of *UBE3A* levels at both time points normalized the decrease in sIPSC frequency (Fig. 5B). *UBE3A* ASO treatments at both 6 and 16 weeks were only partially effective in normalizing the increase in mEPSC frequency (Fig. 5C; differences are not statistically significant), although mEPSC amplitude was corrected by ASO treatment at 6 weeks (Fig. 5C). Moreover, normalizing *UBE3A* levels did not have a significant effect on mIPSC frequency or amplitude (Fig. 5D). To test if the *UBE3A* effect on spontaneous excitatory events is partially caused by changes in spontaneous AP firing, we performed population calcium imaging experiments. Normalizing *UBE3A* at either 6 or 16 weeks reduced the spontaneous frequency of calcium transients in Dup15q neurons (Fig. 5E). These findings suggest that *UBE3A* is necessary for the development of the increased spontaneous excitatory transmission and spontaneous neuronal firing in Dup15q cells. Although normalization of *UBE3A* level at 16 weeks was less effective in reversing some phenotypes compared to treatment at 6 weeks, it successfully normalized the deficit in spontaneous inhibitory transmission (Fig. 5B) and the increase in spontaneous neuronal firing (Fig. 5E).

**Figure 5.**
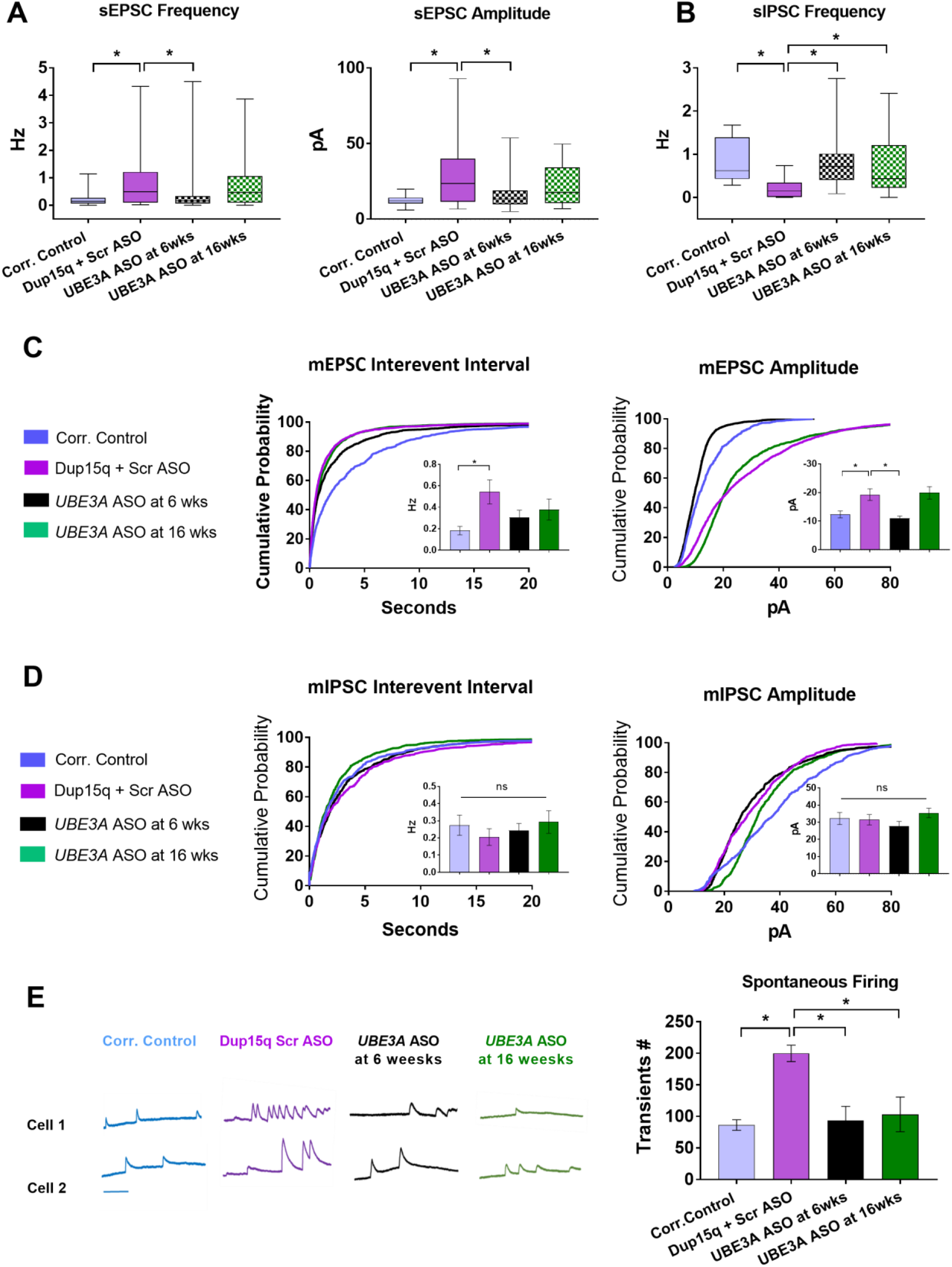
Normalization of *UBE3A* affects synaptic transmission and spontaneous firing phenotypes in Dup15q neurons. **(A)** Frequency (left) and amplitude (right) of spontaneous excitatory postsynaptic currents (sEPSC) in corrected neurons, Dup15q neurons treated with 10 μM scramble ASO, Dup15q neurons treated with 10 μM *UBE3A* ASO at 6 weeks, and Dup15q neurons treated with 10 μM *UBE3A* ASO at 16 weeks (n=22-32 cells per group). **(B)** Frequency of spontaneous inhibitory postsynaptic currents (sIPSC) in corrected neurons, Dup15q neurons treated with 10 μM scramble ASO, Dup15q neurons treated with *UBE3A* ASO at 6 weeks, and Dup15q neurons treated with *UBE3A* ASO at 16 weeks (n= 22-32 cells per group). Box and whisker blots represent the first quartile, median, and third quartile; bars represent 2.5-97.5 percentile outliers. Nonparametric Mann Whitney U test; *, p<0.05. **(C) Left:** Cumulative probability histogram of the interevent interval of all mEPSC events across all cells. **Inset**: Frequency of mEPSC per cell, represented as mean +/- SEM. **Right:** Cumulative probability histogram of the amplitude of all mEPSC events across all cells. **Inset**: Amplitude of mEPSC per cell, represented as mean +/- SEM. (n= 20-29 cells per group). One-way ANOVA and Dunnett’s multiple comparisons tests. **(D) Left:** Cumulative probability histogram of the interevent interval of all mIPSC events across all cells. **Inset**: Frequency of mIPSC per cell, represented as mean +/- SEM. **Right:** Cumulative probability histogram of the amplitude of all mIPSC events across all cells. **Inset**: Amplitude of mIPSCs per cell (n= 15-16 cells per group). One-way ANOVA and Dunnett’s multiple comparisons tests. **(E)** Calcium imaging: **Left:** Example traces of spontaneous calcium transients in two different cells per group. **Right:** Number of spontaneous calcium transient per coverslip over 15 min (n=5 coverslips per group). Scale bar: 100 Seconds. One-way ANOVA and Dunnett’s multiple comparisons tests. *, p< 0.05.

### *UBE3A* overexpression recapitulates key cellular phenotypes in Dup15q neurons

To study the consequences of *UBE3A* overexpression in the context of other duplicated genes in the region, and to confirm that the observed intrinsic excitability and synaptic phenotypes are modulated by *UBE3A*, we increased *UBE3A* dosage in an iPSC line with a paternal duplication in BP2-BP3 of the 11.2 to 13.1 region of chromosome 15 (PatDup). This line, which contains two silenced copies of *UBE3A*, was previously characterized and does not display Dup15q cellular phenotypes (17). *UBE3A* overexpression was achieved by using ASOs targeting *UBE3A-ATS*, a non-coding RNA that silences the paternal copy of *UBE3A*. These ATS-ASOs would theoretically increase *UBE3A* gene dosage from one active copy to three active copies, mimicking the expression level in idic(15) Dup15q syndrome. These cells also have one extra copy of the non-imprinted genes in the 15q11.2-q13 region. We previously confirmed the ability of this specific ASO to reduce *UBE3A*-ATS, leading to an immediate, and long-lasting, increase in *UBE3A* mRNA and protein levels (26).

To determine if early overexpression of *UBE3A* causes the same electrophysiological phenotypes that were observed in Dup15q neurons, ATS ASOs and scramble ASOs (10 µM) were applied to PatDup cells at 6 weeks of *in vitro* development (Experimental design; Fig. 6A). *UBE3A m*RNA was quantified via RT-qPCR and found to be approximately twice the level of *UBE3A* found in scramble ASO-treated PatDup neurons (Fig. 6B). Patch clamp recordings performed at 19 weeks revealed that *UBE3A* overexpression resulted in a hyperexcitability profile similar to that of Dup15q neurons. Specifically, a significant increase in inward current, an increase in AP amplitude, an increase in AP firing frequency, and a hyperpolarized AP threshold were observed (Figs. 6D - 6G). Nevertheless, not all phenotypes were recapitulated by *UBE3A* overexpression. Except for an increase in sIPSC amplitude, neither excitatory nor inhibitory synaptic activity was altered upon *UBE3A* overexpression in PatDup neurons (Fig. 6H, 6I).

**Figure 6.**
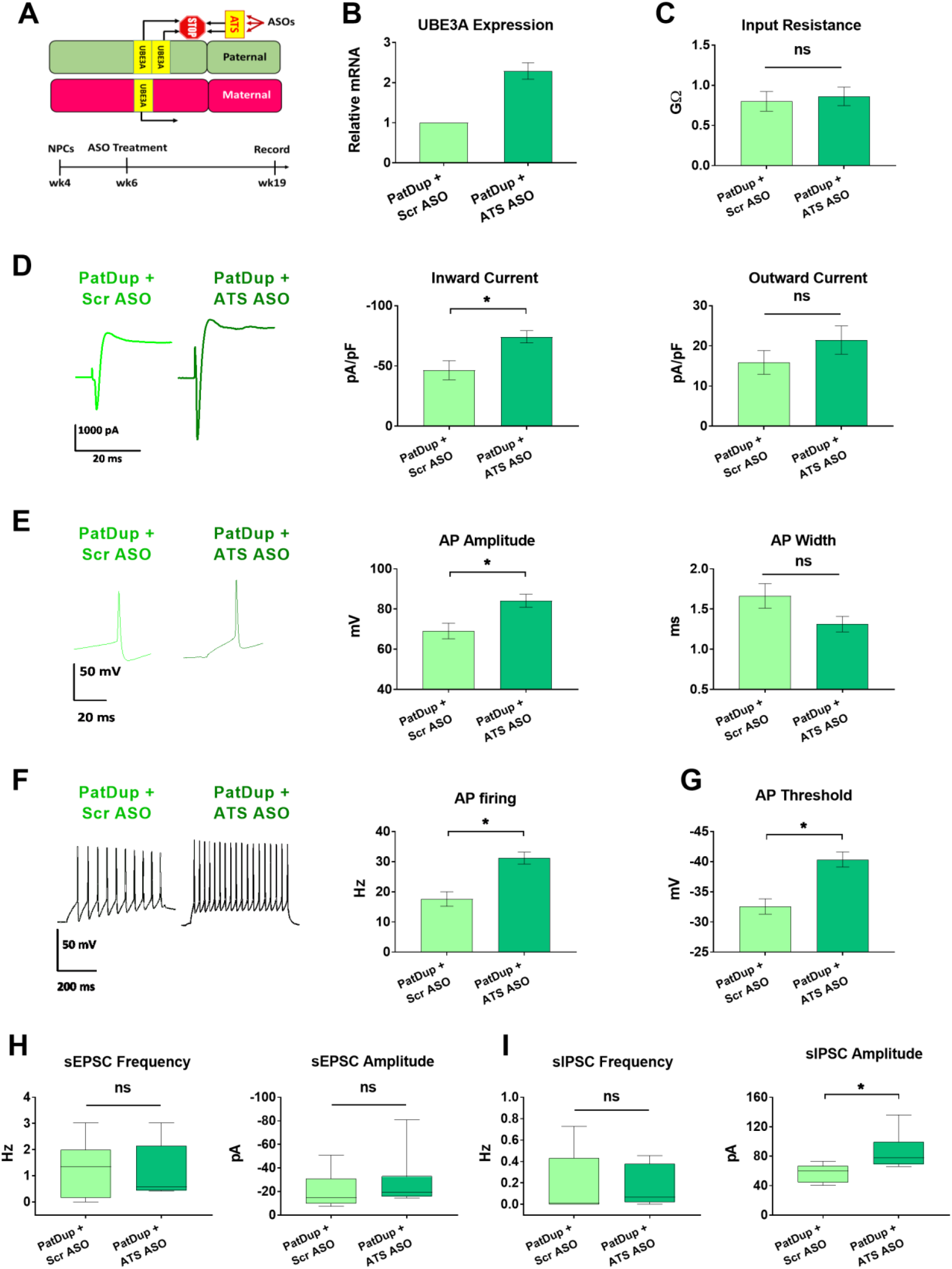
*UBE3A* overexpression recapitulates intrinsic excitability phenotypes but not synaptic deficits of Dup15q neurons. **(A)** Experimental design: Neurons differentiated from an iPSC line with a paternal duplication in Chr.15 encompassing the *UBE3A* region were treated with 10 μM scramble or *UBE3A-ATS* ASOs at 6 weeks. (n = 10 cells per group). **(B)** qPCR showing relative *UBE3A* mRNA expression two weeks post-treatment. **(C)** Input resistance obtained via a 10 mV hyperpolarizing step. Unpaired t-test **(D) Left**: Example inward and outward currents at 19 weeks elicited with a voltage step from -70 mV to +40 mV. **Middle:** Maximum inward current density. **Right:** Maximum outward current density. Student’s t-test. **(E) Left:** Example action potential (AP) traces at 19 weeks elicited by +80 pA current step. **Middle:** Peak AP amplitude. **Right:** AP width at half of the maximum amplitude. Unpaired t-tests. **(F) Left:** Example AP traces showing the firing rate of 19 week-old cells during a 500 ms current step elicited by +80 pA. **Right**: Maximum AP firing rate. Unpaired t-test. **(G)** AP firing threshold. Unpaired t-test. **(H)** Frequency (left) and amplitude (right) of spontaneous excitatory postsynaptic currents (sEPSCs). **(I)** Frequency (left) and amplitude (right) of spontaneous inhibitory postsynaptic currents (sIPSCs). Box and whisker blots represent the first quartile, median, and third quartile; bars represent 2.5-97.5 percentile outliers. Mann Whitney U test; *, p< 0.05.

## Discussion

Maternal duplications in the 11.2-13.1 region of chromosome 15 cause Dup15q syndrome, a disorder characterized by autism, epileptic seizures, and a wide range of intellectual and motor disabilities (1, 27). The parent of origin is important for Dup15q because maternal duplications cause the syndrome, while paternal duplications do not (12, 28, 29). It is thought that *UBE3A*, the gene encoding the ubiquitin protein ligase E3A, plays a critical role in Dup15q phenotypes because it is the only imprinted gene that is expressed solely from the maternal allele in neurons. Animal models of *UBE3A* overexpression have been used to study the syndrome, but with limited success in recapitulating the full range of Dup15q phenotypes (14, 16). Maternal duplication of the syntenic mouse region also failed to replicate behavioral and physiological phenotypes (13), thus, fundamental knowledge regarding the cellular mechanisms of Dup15q and the role of *UBE3A* in the development of the syndrome is lacking.

Here, we employed an innovative CRISPR strategy to successfully eliminate the extra chromosome in an idic(15) Dup15q human iPSC line, thereby creating an isogenic control line. We used these lines to determine the functional differences between Dup15q and corrected neurons and investigate the role of *UBE3A* in Dup15q cellular phenotypes. Dup15q neurons exhibited multiple hyperexcitability phenotypes including an increase in voltage-gated inward current (and the associated increase in AP amplitude), decreased AP width, increased firing, and synaptic changes compared to isogenic corrected control neurons, consistent with previous findings (17).

One critical question for Dup15q is whether a single gene drives the cellular and patient phenotypes. Our data demonstrated that most Dup15q neuronal phenotypes were prevented by normalizing *UBE3A* levels, indicating that excess *UBE3A* is necessary for their development. The majority of idic(15) Dup15q cellular phenotypes were also recapitulated by overexpressing *UBE3A* in PatDup neurons, further supporting a role for excess *UBE3A* in their establishment. Both of these conditions evaluated excess *UBE3A* in the context of duplicated non-imprinted genes. That is, excess *UBE3A* plus excess *GABRB3, GABRA5, GABRG3, HERC2*, and other genes. It is still unknown whether *UBE3A* overexpression alone is sufficient to cause these phenotypes. For instance, deficits in excitatory and inhibitory transmission in Dup15q neurons were not replicated by *UBE3A* overexpression in PatDup neurons, nor fully corrected by normalizing *UBE3A* level in Dup15q neurons. These results support a role for other non-imprinted duplicated genes in the development of synaptic deficits. Nonetheless, even if duplication of the non-imprinted genes contributes to the cellular phenotypes, our data suggest that normalization of *UBE3A* alone may ameliorate the majority of neuronal phenotypes in our human *in vitro* model system. This suggests that therapeutics that target *UBE3A* might influence neuronal excitability and potentially alter the disease course of individuals with Dup15q.

Normalizing *UBE3A* expression at 16 weeks failed to reverse many phenotypes in our experimental paradigm. It is possible that a longer treatment time may be required to reverse these phenotypes in mature neurons, or that optimal connectivity achieved by co-culture with astrocytes or growth as organoids may better support phenotypic rescue. However, there may also be specific therapeutic windows for particular phenotypes. Our iPSC model system is not ideal for teasing out these differences. Interestingly, *UBE3A* normalization at both 6 and 16 weeks rescued the decrease in spontaneous inhibitory synaptic events, and also rescued the increase in spontaneous neuronal firing.

Dup15q is a complicated genetic disorder caused by copy number variation affecting many genes. To begin to dissect the roles of the individual genes, we eliminated the extra chromosome in idic(15) iPSCs to generate the first isogenic human iPSC cell pair. This isogenic pair enabled us to investigate the role of *UBE3A*, a gene thought to play a major role in the disorder, in the development of Dup15q cellular phenotypes. We found that *UBE3A* overexpression is necessary for most of the cellular phenotypes we interrogated. However, some phenotypes - most notably synaptic phenotypes - were not completely dependent on *UBE3A* overexpression, suggesting an important role for non-imprinted genes in the disorder. Future studies to identify the additional gene(s) that contribute to the synaptic alterations in Dup15q neurons may help guide research efforts toward creating a more comprehensive animal model that will better encompass Dup15q behavioral and physiological phenotypes.

Future studies to investigate the mechanism of how excess *UBE3A* and/or other genes cause the neuronal pathophysiology may open the door for the discovery of new therapeutic approaches for Dup15q syndrome.

## Materials and methods

### iPSC lines and CRISPR/Cas9-mediated genome editing

Studies were carried out using iPSC lines generated from two idic(15) Dup15q patients (idic-1 and idic-2) and one individual with a paternal 15q11-q13 interstitial duplication (PatDup). The H9 human embryonic stem cell line and an iPSC line from an unaffected individual were used as controls. These lines were previously characterized (7, 17) and are available from the UConn Cell and Genome Engineering Core. To create an isogenic corrected line, we began with an idic-1 clone that had also been transduced with a fluorescent reporter. Next, sgRNAs targeting GOLGA8, SNORD116, and SNORD115 were designed using MIT’s CRISPR Design Tool (http://crispr.mit.edu; Supplemental Table 2) and cloned into pX459v2.0 (Addgene 62988) vector. Briefly, idic-1 iPSCs were pre-treated with ROCK inhibitor before being dissociated into single cells and nucleofected with CRISPRs. iPSCs were then plated onto puromycin-resistant (DR4) irrMEFs and selected for 48 hours with puromycin (0.5 – 1 μg/ ml). Puromycin-resistant colonies were screened using quantitative PCR-based Copy Number Assays for UBE3A and PML to identify clones that had lost a 15q11-q13 allele and/or a full chromosome 15, respectively. Clones with decreased *UBE3A* copy number, but normal PML copy number were expanded on irrMEFs and then subjected to quantitative DNA methylation analysis at SNRPN to determine the parent-of-origin of the remaining 15q11-q13 alleles. One clone showed a reduction of *UBE3A* and 50% methylation at SNRPN, suggesting that it had lost two maternal copies of chromosome 15q11-q13. This clone was subsequently characterized using a cytoSNP array, metaphase karyotyping, confirmatory quantitative DNA methylation analysis, and gene expression analysis. This characterization revealed no other genetic structural variation compared to the parent line.

### Stem cell culture

Mitomycin C-treated mouse embryonic fibroblasts (Millipore, Burlington MA) and human embryonic stem cell medium (hESC) were used to culture iPSCs. hESC media contained DMEM-F12 (Life Technologies, Carlsbad CA), 0.1 mM non-essential amino acids, 20% knockout serum replacer, 1 mM L-glutamine, 10 ng ml–1 basic fibroblast growth factor, and 0.1 mM 2-mercaptoethanol. A humidified incubator with 5% CO_2_ was used to maintain the cells at 37 °C. Stem cells were mechanically passaged once per week using a 28-gauge needle, and hESC media was replaced daily.

### Neuronal differentiation and maintenance

Neuronal differentiation was carried out according to established monolayer differentiation protocols with minor modifications (7). On day 0, iPSC colonies were cultured for 10 days in N2/B27 neural induction media containing: neurobasal media, 1% N2 supplement, 2% B27 supplement, 2 mM L-glutamine, 1% insulin-transferrin selenium (Life Technologies), 500ng/mL μM of Noggin (R&D Systems, Minneapolis MN) and 2 μM SB431542 (Stemgent, Beltsville MD) were added to the media for 10 days. The media was fully replaced on days 3, 5, 7, 10. On days 11 and 13, neurons were fed with N2/B27 media without Noggin or SB431542. On day 14, neuronal rosettes were passaged as small clusters in a 1:2 ratio using EZPassage (Invitrogen, Waltham MA) and plated on poly-O/laminin-coated 6-well plates. Fifty percent media replacement was carried out on days 17 and 19. On day 21, accutase (Millipore) was used to dissociate the cells. Cells were then counted using a hemocytometer and plated on polyornithine/laminin-coated 6-well plates at a density of 1 × 10^6^ cells/well with 10 μM ROCK inhibitor Y27632 (Wako, Richmond VA). Fifty percent media replacement was carried out on days 24 and 26. On day 28, Accutase was used to dissociate the cells, and NPCs (neural progenitor cells) were counted using a hemocytometer, resuspended in NPC freezing media (1 × 10^6^ cells/vial) containing 90% fetal bovine serum (Invitrogen) and 10% DMSO (Sigma-Aldrich, St. Louis MO). Vials were stored at -80^0^ C for 24 hours, and transferred to liquid nitrogen for long-term storage. To perform experiments, neurons were thawed in neural differentiation medium (NDM) containing growth factors (Neurobasal, 1 × B-27 supplement, 0.1 mM NEAA, 2 mM L-glutamine, 1 μg/ml laminin, 1 μM cAMP (Sigma), 0.2 mM L-ascorbic acid (Sigma), 20 ng/ml glial cell line-derived neurotrophic factor and 20 ng/ml brain-derived neurotrophic factor (Peprotech, East Windsor NJ). Cells were plated on polyornithine/laminin-coated wells. After one week, NPCs were dissociated using accutase, counted using a hemocytometer, and plated on polyornithine/laminin-coated glass coverslips at a density of 20,000 cells/coverslip. To aid in cell attachment, ROCK inhibitor Y-27632 (10 μM; Wako) was added to the NDM media during initial plating. Cells were maintained with no antibiotics in NDM media, and 50% media replacement was carried out twice a week. A minimum of two independent rounds of neuronal differentiations were carried out for each cell line.

### Antisense oligonucleotides (ASOs)

*UBE3A* ASOs and *UBE3A-ATS* ASOs were used to knock down or overexpress *UBE3A*, respectively. Control scramble sequence ASOs were also used. All ASOs were provided by Ionis Pharmaceuticals (Carlsbad CA) and synthesized as previously described (24). ASOs were 20 bp in length with ten DNA nucleotides in the center, a phosphorothioate backbone, and five 2′-O-methoxyethyl-modified nucleotides at each end. The ASOs were added directly to the culture media at 10 μM concentration. After 72 hours, the media was completely removed and cells were fed with fresh media. iPSC-derived neurons exposed to a single treatment with *UBE3A* ASOs led to sustained mRNA and protein knockdown (25, 26). The sequences of all ASOs used in the study are provided in Supplemental Table 3.

### Electrophysiology

Coverslips were transferred to a recording chamber fixed to an Olympus BX51WI upright microscope stage. Neurons were visualized using a 40x water-immersion lens and identified based on morphology. A continuous flow of oxygenated artificial cerebrospinal fluid (aCSF) containing (mM): 125 NaCl, 2.5 KCl, 15.0 dextrose, 1.25 NaH_2_PO_4_, 2.0 MgCl_2_-6H_2_O, 25.0 NaHCO_3_, and 2.0 CaCl_2_ was maintained at a rate of 1.5 ml/min. All recordings were done at room temperature.

Pipettes with a resistance ranging from 5-8 MΩ were pulled from borosilicate glass capillaries using Flaming/Brown P-97 micropipette puller and filled with an internal solution containing (mM): 125.0 K-gluconate, 4.0 KCl, 10.0 HEPES, 10.0 phosphocreatine, 4.0 Na2-ATP, 0.3 Na-GTP, 0.20 CaCl2, and 1.0 EGTA. Input resistance was monitored throughout the recordings by applying a 10 mV hyperpolarizing step from -70 mV. Resting membrane potential (RMP) was noted in current-clamp mode by injecting 0 current. RMP values were corrected for the liquid junction potential calculated offline (JPcalc). Inward and transient outward currents were recorded in voltage-clamp mode by holding the cells at -70 mV and applying 300 ms depolarizing steps from -100 to +40 mV in 10 mV increments. AP firing was elicited in current-clamp mode by injecting current to hold the cells at ∼-70 mV and applying 500 ms current steps from -10 to +80 pA in 10 pA increments. Spontaneous excitatory synaptic currents were monitored in voltage-clamp mode at a holding potential of -70 mV, and spontaneous inhibitory synaptic currents were monitored at a holding potential of 0 mV. Miniature synaptic events were recorded using the same protocols in the presence of 1 μM TTX. Electrophysiological analysis was performed using Clampfit (Molecular Devices, San Jose CA).

### Calcium imaging

X-Rhod-1 dye (Thermofisher), reconstituted with DMSO, was added to the cell culture media for a final concentration of 2 μM. Coverslips were incubated with this mixture for 1 hour at 37°C then transferred to the recording chamber and perfused with aCSF for 15 min before imaging. Neurons were imaged using a Cairn OptoLED light source system at 594 nM (Texas Red), SM-CCD67 camera at 100 Hz, and Turbo-SM software. Analysis of the data was performed as previously described using Fluorescence Single Neuron and Network Analysis Package (FluoroSNNAP) (25). Spontaneous activity was recorded for 15 min from each coverslip.

### Immunocytochemistry

To fix the neurons, coverslips were incubated for 15 min in 4% paraformaldehyde in phosphate-buffered saline (PBS), and washed three times with PBS (5 min each). Neurons were permeabilized for 10 minutes at room temperature using a permeabilization buffer containing 0.2% Triton X-100 and a blocking buffer. Next, a blocking buffer containing 10% normal goat serum, 1% BSA, 0.3M glycine, and 0.2% Tween 20 diluted in 1x PBS was used to block for non-specific binding for 1 hour at room temperature. Primary antibodies (PSD95 1:500, Antibodies Inc., Davis CA and MAP2 1:1000; Millipore) were diluted in 1x PBS with 1% BSA and 0.2% Tween. Coverslips were incubated overnight at 4°C, then washed three times with 1x PBS with 0.2% Tween 20 for 5 min each. Secondary antibodies (Alexa Fluor; 594 Goat Anti-Rabbit and 647 Goat Anti-Mouse) were diluted in 1x PBS with 1% BSA and 0.2% Tween 20. Coverslips were incubated in the dark for 1 hour at room temperature, then washed three times in 1x PBS with 0.2% Tween 20 and mounted on slides using hard set Vectashield mounting medium containing DAPI stain (Vector Laboratories, Burlingame CA). A Zeiss Axiovision fluorescence microscope was used for imaging. For quantification, a minimum of 4-5 images were captured per coverslip. Process length was traced and measured with ImageJ, and the number of PSD95 puncta along MAP2 processes was counted by an individual blinded to the experimental conditions.

### qPCR

RNA was extracted with RNAeasy extraction kit (Qiagen, Hilden Germany) following the manufacturer’s protocol. High-capacity cDNA Reverse Transcription kit was used to generate cDNA (Applied Biosystems, Waltham MA). Quantitative real-time PCR for *UBE3A* was performed using TaqMan gene expression assays (Applied Biosystems) and CFX Connect Real-Time System (Bio-Rad Laboratories, Hercules CA). The expression level was normalized to the housekeeping gene GAPDH (Applied Biosystems).

### Statistical Analysis

Prism software (GraphPad, San Diego CA) was used for all statistical analysis. For normally distributed data, parametric Student’s t-test or one-way repeated measures analysis of variance (ANOVA) and Dunnett’s multiple comparisons tests were used as indicated in the respective graphs. Data are presented as mean +/- standard error of the mean. For data that are not normally distributed, a non-parametric Mann Whitney U test was used, and data were represented in box and whisker blots representing the median, first, and third quartiles, and error bars representing 2.5-97.5 percentile outliers. All statistical significance values of less than 0.05 are represented by an asterisk.

## Acknowledgments

This work was supported by NIH Grants NS111965 (ESL, SJC) and NS111986 (ESL), the Eagles Autism Foundation (ESL), and the Schlumberger Foundation (ME).

## Author contributions

ME, CC, and DG conducted experiments and analyzed data. AD and CS carried out genome editing to generate the isogenic cell line. TMR designed and optimized experimental procedures. ME, SJC, and ESL designed the project and wrote the manuscript. All authors were involved in the preparation and review of the manuscript and approved the final submitted version.

## References

1. DiStefano C, Gulsrud A, Huberty S, Kasari C, Cook E, Reiter LT, Thibert R, and Jeste SS. Identification of a distinct developmental and behavioral profile in children with Dup15q syndrome. J Neurodev Disord. 2016;8(19.

2. Al Ageeli E, Drunat S, Delanoe C, Perrin L, Baumann C, Capri Y, Fabre-Teste J, Aboura A, Dupont C, Auvin S, et al. Duplication of the 15q11-q13 region: clinical and genetic study of 30 new cases. Eur J Med Genet. 2014;57(1):5–14.

3. Urraca N, Cleary J, Brewer V, Pivnick EK, McVicar K, Thibert RL, Schanen NC, Esmer C, Lamport D, and Reiter LT. The interstitial duplication 15q11.2-q13 syndrome includes autism, mild facial anomalies and a characteristic EEG signature. Autism Res. 2013;6(4):268–79.

4. Conant KD, Finucane B, Cleary N, Martin A, Muss C, Delany M, Murphy EK, Rabe O, Luchsinger K, Spence SJ, et al. A survey of seizures and current treatments in 15q duplication syndrome. Epilepsia. 2014;55(3):396–402.

5. Friedman D, Thaler A, Thaler J, Rai S, Cook E, Schanen C, and Devinsky O. Mortality in isodicentric chromosome 15 syndrome: The role of SUDEP. Epilepsy Behav. 2016;61(1–5.

6. DiStefano C, Wilson RB, Hyde C, Cook EH, Thibert RL, Reiter LT, Vogel-Farley V, Hipp J, and Jeste S. Behavioral characterization of dup15q syndrome: Toward meaningful endpoints for clinical trials. Am J Med Genet A. 2020;182(1):71–84.

7. Germain ND, Chen PF, Plocik AM, Glatt-Deeley H, Brown J, Fink JJ, Bolduc KA, Robinson TM, Levine ES, Reiter LT, et al. Gene expression analysis of human induced pluripotent stem cell-derived neurons carrying copy number variants of chromosome 15q11-q13.1. Mol Autism. 2014;5(44.

8. Huang L, Kinnucan E, Wang G, Beaudenon S, Howley PM, Huibregtse JM, and Pavletich NP. Structure of an E6AP-UbcH7 complex: insights into ubiquitination by the E2-E3 enzyme cascade. Science. 1999;286(5443):1321–6.

9. Nawaz Z, Lonard DM, Smith CL, Lev-Lehman E, Tsai SY, Tsai MJ, and O’Malley BW. The Angelman syndrome-associated protein, E6-AP, is a coactivator for the nuclear hormone receptor superfamily. Mol Cell Biol. 1999;19(2):1182–9.

10. Low D, and Chen KS. Genome-wide gene expression profiling of the Angelman syndrome mice with Ube3a mutation. Eur J Hum Genet. 2010;18(11):1228–35.

11. Chamberlain SJ, and Lalande M. Neurodevelopmental disorders involving genomic imprinting at human chromosome 15q11-q13. Neurobiol Dis. 2010;39(1):13–20.

12. Cook EH, Jr., Lindgren V, Leventhal BL, Courchesne R, Lincoln A, Shulman C, Lord C, and Courchesne E. Autism or atypical autism in maternally but not paternally derived proximal 15q duplication. Am J Hum Genet. 1997;60(4):928–34.

13. Nakatani J, Tamada K, Hatanaka F, Ise S, Ohta H, Inoue K, Tomonaga S, Watanabe Y, Chung YJ, Banerjee R, et al. Abnormal behavior in a chromosome-engineered mouse model for human 15q11-13 duplication seen in autism. Cell. 2009;137(7):1235–46.

14. Smith SE, Zhou YD, Zhang G, Jin Z, Stoppel DC, and Anderson MP. Increased gene dosage of Ube3a results in autism traits and decreased glutamate synaptic transmission in mice. Sci Transl Med. 2011;3(103):103ra97.

15. Krishnan V, Stoppel DC, Nong Y, Johnson MA, Nadler MJ, Ozkaynak E, Teng BL, Nagakura I, Mohammad F, Silva MA, et al. Autism gene Ube3a and seizures impair sociability by repressing VTA Cbln1. Nature. 2017;543(7646):507–12.

16. Copping NA, Christian SGB, Ritter DJ, Islam MS, Buscher N, Zolkowska D, Pride MC, Berg EL, LaSalle JM, Ellegood J, et al. Neuronal overexpression of Ube3a isoform 2 causes behavioral impairments and neuroanatomical pathology relevant to 15q11.2-q13.3 duplication syndrome. Hum Mol Genet. 2017;26(20):3995–4010.

17. Fink JJ, Schreiner JD, Bloom JE, James J, Baker DS, Robinson TM, Lieberman R, Loew LM, Chamberlain SJ, and Levine ES. Hyperexcitable Phenotypes in Induced Pluripotent Stem Cell-Derived Neurons From Patients With 15q11-q13 Duplication Syndrome, a Genetic Form of Autism. Biol Psychiatry. 2021;90(11):756–65.

18. Gupta N, Susa K, Yoda Y, Bonventre JV, Valerius MT, and Morizane R. CRISPR/Cas9-based Targeted Genome Editing for the Development of Monogenic Diseases Models with Human Pluripotent Stem Cells. Curr Protoc Stem Cell Biol. 2018;45(1):e50.

19. Wang H, Yang H, Shivalila CS, Dawlaty MM, Cheng AW, Zhang F, and Jaenisch R. One-step generation of mice carrying mutations in multiple genes by CRISPR/Cas-mediated genome engineering. Cell. 2013;153(4):910–8.

20. Xiao A, Wang Z, Hu Y, Wu Y, Luo Z, Yang Z, Zu Y, Li W, Huang P, Tong X, et al. Chromosomal deletions and inversions mediated by TALENs and CRISPR/Cas in zebrafish. Nucleic Acids Res. 2013;41(14):e141.

21. Adikusuma F, Williams N, Grutzner F, Hughes J, and Thomas P. Targeted Deletion of an Entire Chromosome Using CRISPR/Cas9. Mol Ther. 2017;25(8):1736–8.

22. Zuo E, Huo X, Yao X, Hu X, Sun Y, Yin J, He B, Wang X, Shi L, Ping J, et al. CRISPR/Cas9-mediated targeted chromosome elimination. Genome Biol. 2017;18(1):224.

23. DeVos SL, and Miller TM. Antisense oligonucleotides: treating neurodegeneration at the level of RNA. Neurotherapeutics. 2013;10(3):486–97.

24. Meng L, Ward AJ, Chun S, Bennett CF, Beaudet AL, and Rigo F. Towards a therapy for Angelman syndrome by targeting a long non-coding RNA. Nature. 2015;518(7539):409–12.

25. Fink JJ, Robinson TM, Germain ND, Sirois CL, Bolduc KA, Ward AJ, Rigo F, Chamberlain SJ, and Levine ES. Disrupted neuronal maturation in Angelman syndrome-derived induced pluripotent stem cells. Nat Commun. 2017;8(15038.

26. Germain ND, Gorka D, Drennan R, Jafar-nejad P, Whipple A, Core L, Levine ES, Rigo F, and Chamberlain SJ. Antisense oligonucleotides targeting UBE3A-ATS restore expression of UBE3A by relieving transcriptional interference. bioRxiv. 2021:2021.07.09.451826.

27. Kalsner L, and Chamberlain SJ. Prader-Willi, Angelman, and 15q11-q13 Duplication Syndromes. Pediatr Clin North Am. 2015;62(3):587–606.

28. Mao R, Jalal SM, Snow K, Michels VV, Szabo SM, and Babovic-Vuksanovic D. Characteristics of two cases with dup(15)(q11.2-q12): one of maternal and one of paternal origin. Genet Med. 2000;2(2):131–5.

29. Marini C, Cecconi A, Contini E, Pantaleo M, Metitieri T, Guarducci S, Giglio S, Guerrini R, and Genuardi M. Clinical and genetic study of a family with a paternally inherited 15q11-q13 duplication. Am J Med Genet A. 2013;161A(6):1459–64.

