## Supplemental Data for "The role of *UBE3A* in the autism and epilepsy-related Dup15q syndrome using patient-derived, CRISPR-corrected neurons"

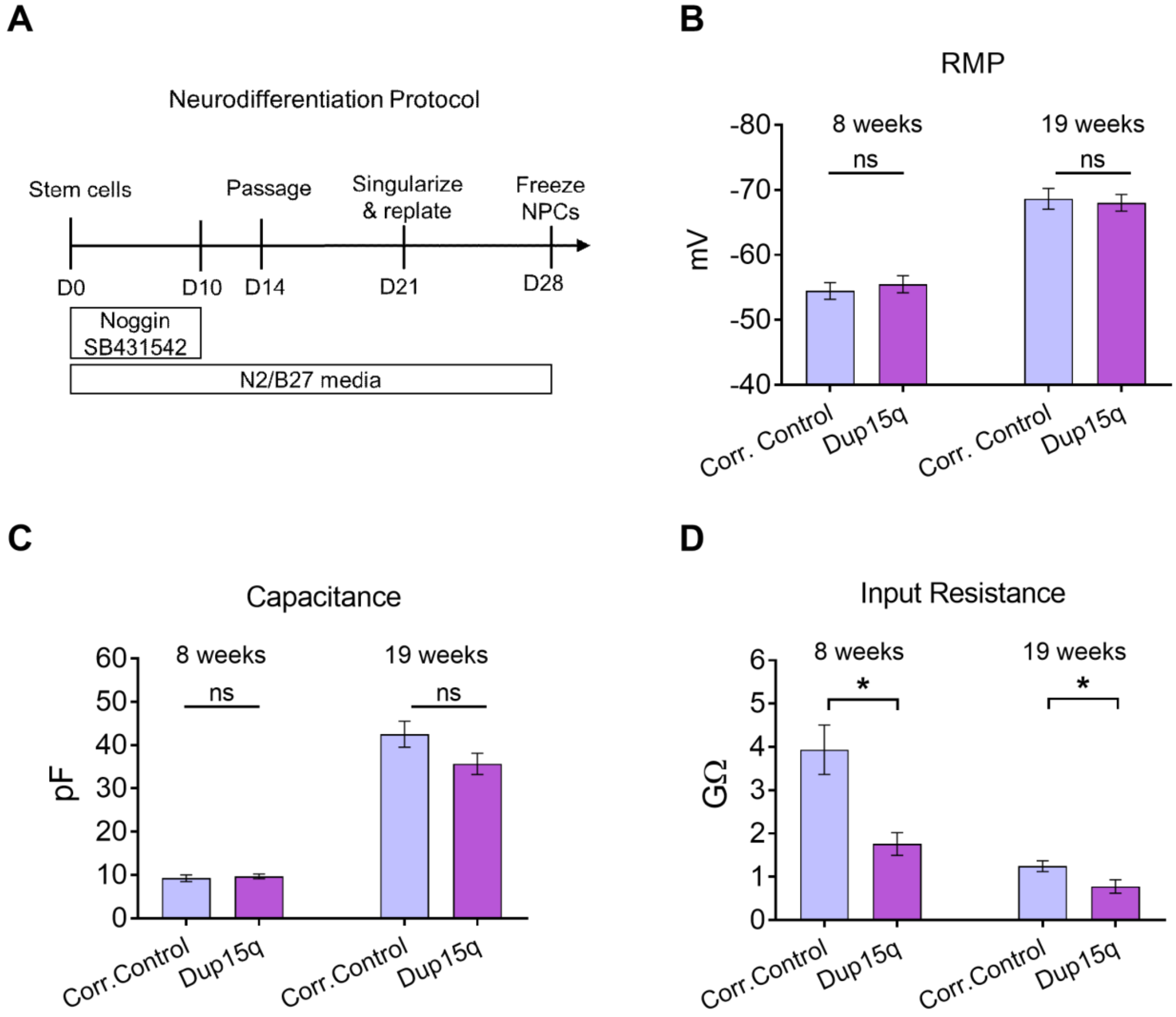

**Supplemental Fig. 1: Neural differentiation protocol and additional electrophysiological properties of Dup15q neurons. (A)** Neural differentiation protocol: modified dual-SMAD inhibition protocol using Noggin and SB431542 for 10 days. **(B)** Resting membrane potential (RMP) of Dup15q neurons and corrected controls at 8 and 19 weeks *in vitro*. **(C)** Cell capacitance at 8 and 19 weeks. **(D)** Input resistance at 8 and 19 weeks. (n= 20-25 cells in all groups). Unpaired t-tests; *, $p < 0.05$.

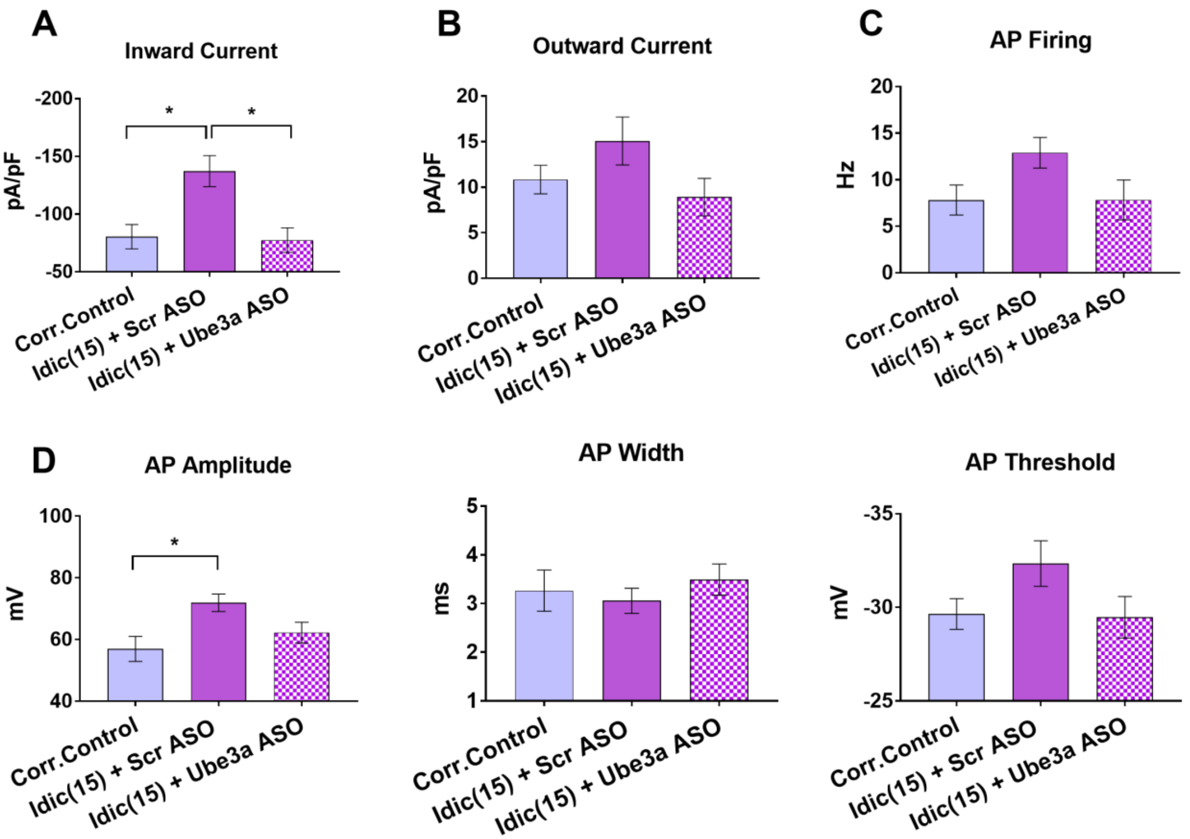

**Supplemental Fig. 2: Effects of *UBE3A* normalization at 8 weeks of *in vitro* development.**

Dup15q neurons were treated with either scramble ASO or *UBE3A* ASO at 6 weeks, and patch-clamp recordings were performed at 8 weeks of development (n=22-32 cells/group). **(A)** Maximum inward current density. **(B)** Maximum outward current density. **(C)** Maximum firing rate during 500 ms current steps from -10 to +80 pA. **(D)** Action potential (AP) characteristics. **Left:** AP peak amplitude, **Middle:** AP width measured at half maximum amplitude; **Right:** AP firing threshold. One-way ANOVA and Dunnett's multiple comparisons tests were performed. *, $p < 0.05$.

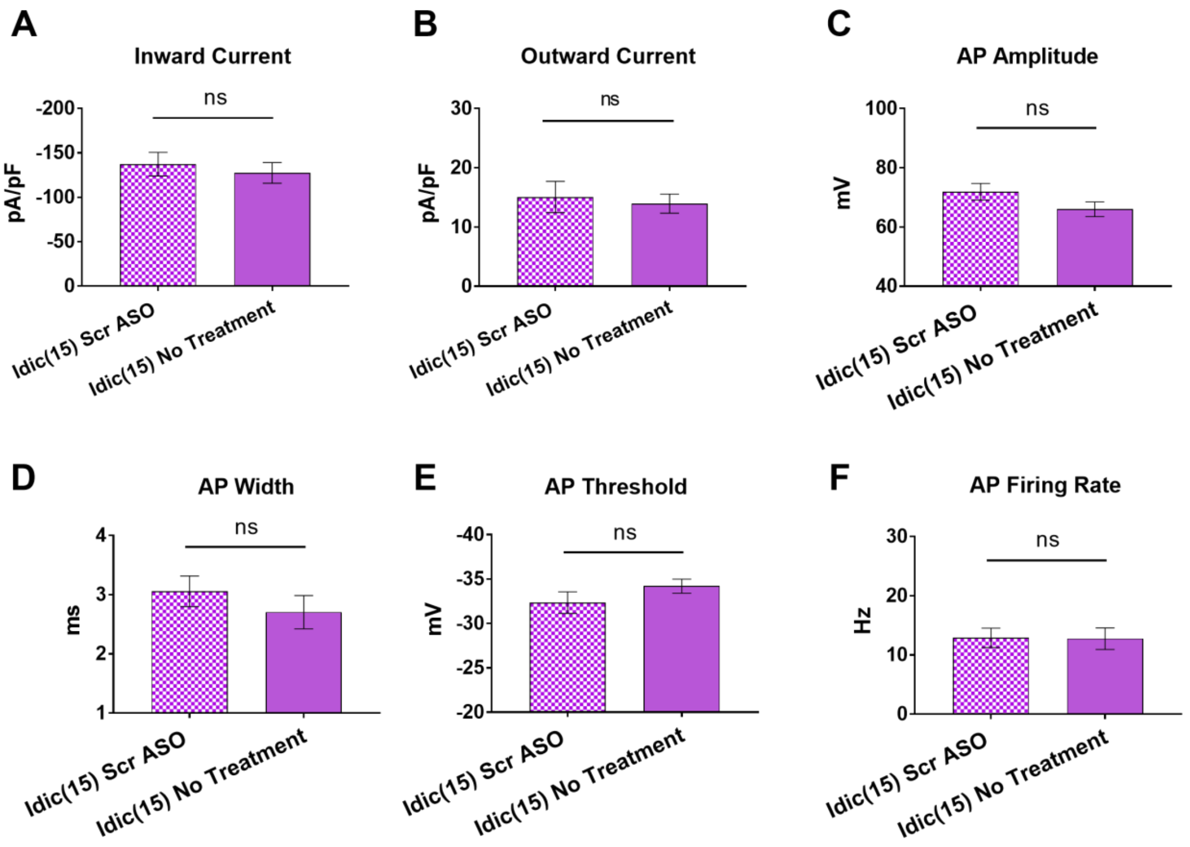

**Supplemental Fig. 3: Scramble control ASO does not affect neuronal function.** Scramble ASO was added to Dup15q neurons at 6 weeks of development, and patch-clamp recordings were performed at 8 weeks. No differences were found between treated and untreated cells. **(A)** inward current density, **(B)** outward current density, **(C)** action potential (AP) amplitude, **(D)** AP width, **(E)** AP threshold, and **(F)** AP firing rate. (n= 20-24 cells in all groups). Unpaired t-tests.

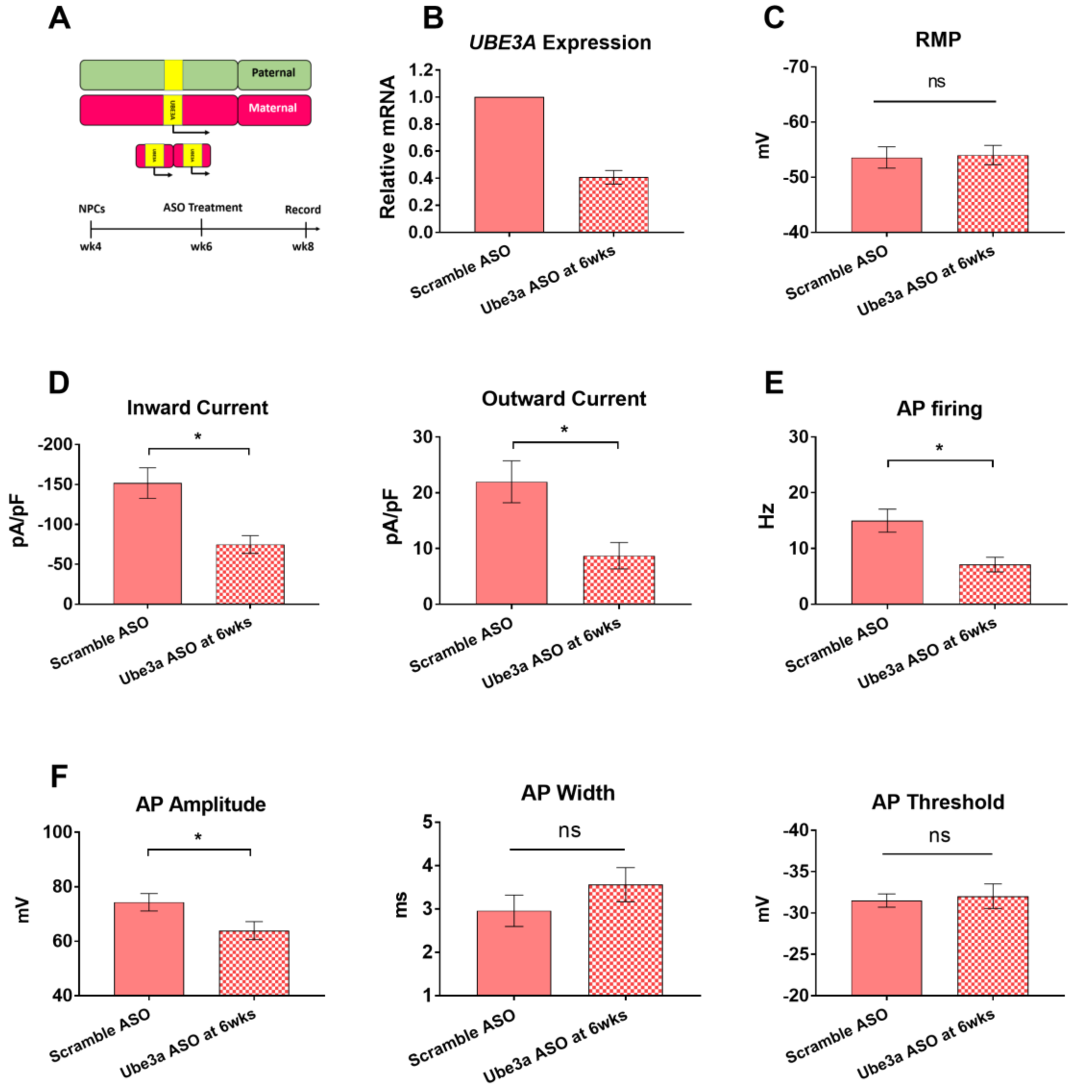

**Supplemental Fig. 4: Normalization of *UBE3A* level prevents intrinsic excitability phenotypes in a second idic(15) Dup15q line.** Dup15q cells were treated with either scramble ASO or *UBE3A* ASO at 6 weeks, and patch-clamp recordings were performed at 8 weeks (n=16 cells per group). **(A)** Chromosomal illustration and experimental design. **(B)** qPCR showing *UBE3A* mRNA relative expression 2 weeks post-ASO treatment. **(C)** Resting membrane potential. **(D)** Maximum inward (left) and outward (right) currents normalized to cell capacitance. **(E)** Maximum firing rate during 500 ms current steps from -10 to +80 pA. **(F)** Action potential (AP) characteristics. **Left:** AP peak amplitude, **Middle:** AP width measured at half maximum amplitude; **Right:** AP firing threshold. Unpaired t-tests; *, $p < 0.05$.

| gRNA name | sequence |
|---|---|
| GOLGA8 | CTGGGTGTGAGGGCACGTGG |
| SNORD116 | CATTTTGTTCAGCTTTTCCA |
| SNORD115 | TGCTCAATAGGATTATGCTG |

**Supplemental Table 1. CRISPR guide RNAs used for generation of corrected Dup iPSC line.**

| Idic-1 CytoSNP Results | | | | | |
|---|---|---|---|---|---|
| ISCN | Type | Chromosome | Start | End | Impact |
| 5p15.33-5p15.33 | GAIN | 5 | 2,981,865 | 3,454,482 | Gain of 473Kb (<1Mb), and overlaps 1 HGNC and 0 OMIM gene(s). |
| 5q22.1-5q23.1 | LOSS | 5 | 109,817,549 | 118,073,885 | Loss of 8256Kb (>=1Mb), and overlaps 56 HGNC and 26 OMIM gene(s). It overlaps the known disease region(s): OMIM disease: ADENOMATOUS POLYPOSIS OF THE COLON; APC |
| 11p15.4-11p15.4 | LOSS | 11 | 9,275,898 | 9,776,567 | Loss of 501Kb (<1Mb), and overlaps 9 HGNC and 4 OMIM gene(s). |
| 15q11.1-15q13.3 | GAIN | 15 | 20,071,673 | 32,514,341 | Gain of 12,442Kb (>=1Mb), and overlaps 245 HGNC and 38 OMIM gene(s). It overlaps the known disease region(s): OMIM disease: PRADER-WILLI SYNDROME; PWS (176270), OMIM disease: ANGELMAN SYNDROME; AS, OMIM disease: CHROMOSOME 15q13.3 DELETION SYNDROME |
| Xq13.1-Xq21.1 | LOH | X | 71,406,301 | 77,848,134 | LOH region of 6442Kb (>=1Mb), and overlaps 89 HGNC and 29 OMIM gene(s). It overlaps the known disease region(s): OMIM disease: X INACTIVATION-SPECIFIC TRANSCRIPT; XIST |
| **Corrected Control CytoSNP Results** | | | | | |
| ISCN | Type | Chromosome | Start | End | Impact |
| 5p15.33-5p15.33 | GAIN | 5 | 2,972,677 | 3,454,482 | Gain of 482Kb (<1Mb), and overlaps 1 HGNC and 0 OMIM gene(s). |
| 5q22.1-5q23.1 | LOSS | 5 | 109,817,549 | 118,082,030 | Loss of 8264Kb (>=1Mb), and overlaps 56 HGNC and 26 OMIM gene(s). It overlaps the known disease region(s): OMIM disease: ADENOMATOUS POLYPOSIS OF THE COLON; APC |
| 11p15.4-11p15.4 | LOSS | 11 | 9,275,898 | 9,776,567 | Loss of 501Kb (<1Mb), and overlaps 9 HGNC and 4 OMIM gene(s). |
| Xq13.1-Xq21.1 | LOH | X | 71,406,301 | 77,848,134 | LOH region of 6442Kb (>=1Mb), and overlaps 89 HGNC and 29 OMIM gene(s). It overlaps the known disease region(s): OMIM disease: X INACTIVATION-SPECIFIC TRANSCRIPT; XIST |

**Supplemental Table 2. CytoSNP analysis of idic-1 and isogenic corrected control lines.**

| ASO name | sequence |
|---|---|
| UBE3A ASO | 5′-TGAGCTATCACCTATCCTTG-3′ |
| UBE3A-ATS ASO | 5′-AGTAAGGTCTGTTATTCTCC -3′ |
| Scramble ASO | 5′-CCTTCCCTGAAGGTTCCTCC-3′ |

**Supplemental Table 3. Antisense oligonucleotide sequences.**
